# Subspace partitioning in human prefrontal cortex resolves cognitive interference

**DOI:** 10.1101/2022.11.16.516719

**Authors:** Jan Weber, Gabriela Iwama, Anne-Kristin Solbakk, Alejandro O. Blenkmann, Pal G. Larsson, Jugoslav Ivanovic, Robert T. Knight, Tor Endestad, Randolph Helfrich

## Abstract

Human prefrontal cortex (PFC) constitutes the structural basis underlying flexible cognitive control, where mixed-selective neural populations encode multiple task-features to guide subsequent behavior. The mechanisms by which the brain simultaneously encodes multiple task-relevant variables while minimizing interference from task-irrelevant features remain unknown. Leveraging intracranial recordings from the human PFC, we first demonstrate that competition between co-existing representations of past and present task variables incurs a behavioral switch cost. Our results reveal that this interference between past and present states in the PFC is resolved through coding partitioning into distinct low-dimensional neural states; thereby strongly attenuating behavioral switch costs. In sum, these findings uncover a fundamental coding mechanism that constitutes a central building block of flexible cognitive control.

## Introduction

In a complex and ever-changing environment, humans need to dynamically adjust their actions according to immediate contextual demands. For example, switching between braking and accelerating in a traffic jam requires rapid shifts between competing actions. The ability to flexibly adjust in an ever-changing environment hallmarks flexible cognitive control. However, rapid shifts between competing actions often incur a behavioral cost, i.e. slower and more error-prone responses following a shift from one action to another.

Multiple lines of research in humans and animal models provide converging evidence that the prefrontal cortex (PFC) constitutes the key structure that enables cognitive flexibility to guide adaptive goal-directed behavior^1-3^. To support flexible operations, PFC neurons are not feature-specific, but instead exhibit mixed selectivity and context-dependent coding of task features (i.e. sensory stimuli, abstract rules or actions)^4-7^. Mixed-selective neurons contribute to multiple cognitive operations by participating in different transient coalitions of cell assemblies. Consequently, information encoded by mixed-selective neurons can only be understood at the level of population activity. This notion is now referred to as the population doctrine, which posits that neural populations reflect the fundamental unit of computation in the brain^8,9^. The population doctrine further postulates that rapid switches between distinct cognitive operations can efficiently be implemented by adjusting the population geometry, i.e. the re-configuration of cell assemblies. A key advantage of this model is that neural representations that evolve in parallel can be integrated into a unified, conjunctive representation^10-12^. In support of this hypothesis, recent findings demonstrated that conjunctive coding schemes enable the flexible context-dependent remapping between sensory inputs and behavioral outputs^12^; predicting behavioral performance on a trial-by-trial basis^10^.

However, parallel encoding of different variables implies that not all encoded features are behaviorally-relevant, hence, raising the question if and how encoding of task-irrelevant (latent) factors impacts subsequent behavior. Previously, it had been observed that past choices reflect typical latent factors, which impact current task-relevant representations^13-16^. At the behavioral level previous choices modulate subsequent behavior (i.e. serial response bias^13,16-18^). Integration of knowledge about past choices is oftentimes desirable to correct past mistakes^19^, but might cause interference when consecutive choices are independent, thus, giving rise to behavioral switch costs^20-22^. In this scenario, a conjunctive neural code that integrates information about the past is detrimental for behavioral performance^15^. To date, little is known about how the human brain integrates multiple task-relevant representations, while minimizing interference from task-irrelevant, latent factors, in order to guide subsequent behavior. Theoretical work has proposed that neural populations can reduce interference by orthogonalizing competing representations^23,24^. This change in population geometry enables downstream regions to flexibly readout information about competing states from the same neural population^12^. Evidence in support of this notion stems from a recent study in mouse auditory cortex that demonstrated that past and present sensory representations reside in orthogonal subspaces^25^. However, it remained unaddressed if the identified neural representations had immediate behavioral relevance. Hence, to date it remains unknown if similar principles also apply to the human brain, especially in higher-order association cortex and whether orthogonalization constitutes a key mechanism to reduce interference between competing neural representations^26^.

Here, we addressed these outstanding questions by directly recording intracranial electroencephalography (iEEG) from the human prefrontal and motor cortex while participants performed a modified stop-signal task that enabled disentangling how competing neural representations between past states and current goals influence human decision-making. We leveraged the power of analyzing high-frequency activity (HFA) as a direct approximation of local neural population activity^27-29^. We specifically tested if neural population activity in PFC and motor cortex simultaneously encode information about the past and present. The key question was whether overlapping neural representations account for behavioral switch costs between movement inhibition and execution. We hypothesized that efficient distributed computing at the level of population activity constituted a core mechanism to avoid interference between distinct latent factors. Collectively, we tested whether efficient cortical partitioning of distinct neural representations enables flexible cognitive control.

## Results

We recorded intracranial EEG (iEEG) from 19 pharmaco-resistant patients with epilepsy (33.73 years ± 12.52, mean ± SD; 7 females) who performed a predictive motor task (**Fig. 1a**). Participants were instructed to closely track a vertically moving target and respond as soon as the moving target reached a predefined spatial location (*go-trials*). In a subset of trials, the moving target stopped prematurely and participants were instructed to withhold their response (*stop-trials*). A contextual cue indicated the likelihood of a premature stop (referred to as *predictive context; Methods)*. Previously, we demonstrated that the human PFC integrates current contextual information to guide goal-directed behavior^30^. Additionally, human behavior is also strongly modulated by a variety of latent factors, such as the tendency to systematically repeat choices, known as history-dependent serial biases^13,17,31^. We here investigated the neural mechanisms underlying such history-dependent serial biases, quantified as the behavioral switch costs between distinct action demands – initiation or inhibition of a goal-directed movement. Thus, task-switching was defined based on the congruency between the trial type (*stop/go-trial*) at trial *n* and *n-1* (*congruent = go-trial followed by go-trial; incongruent = stop-trial followed by go-trial*).

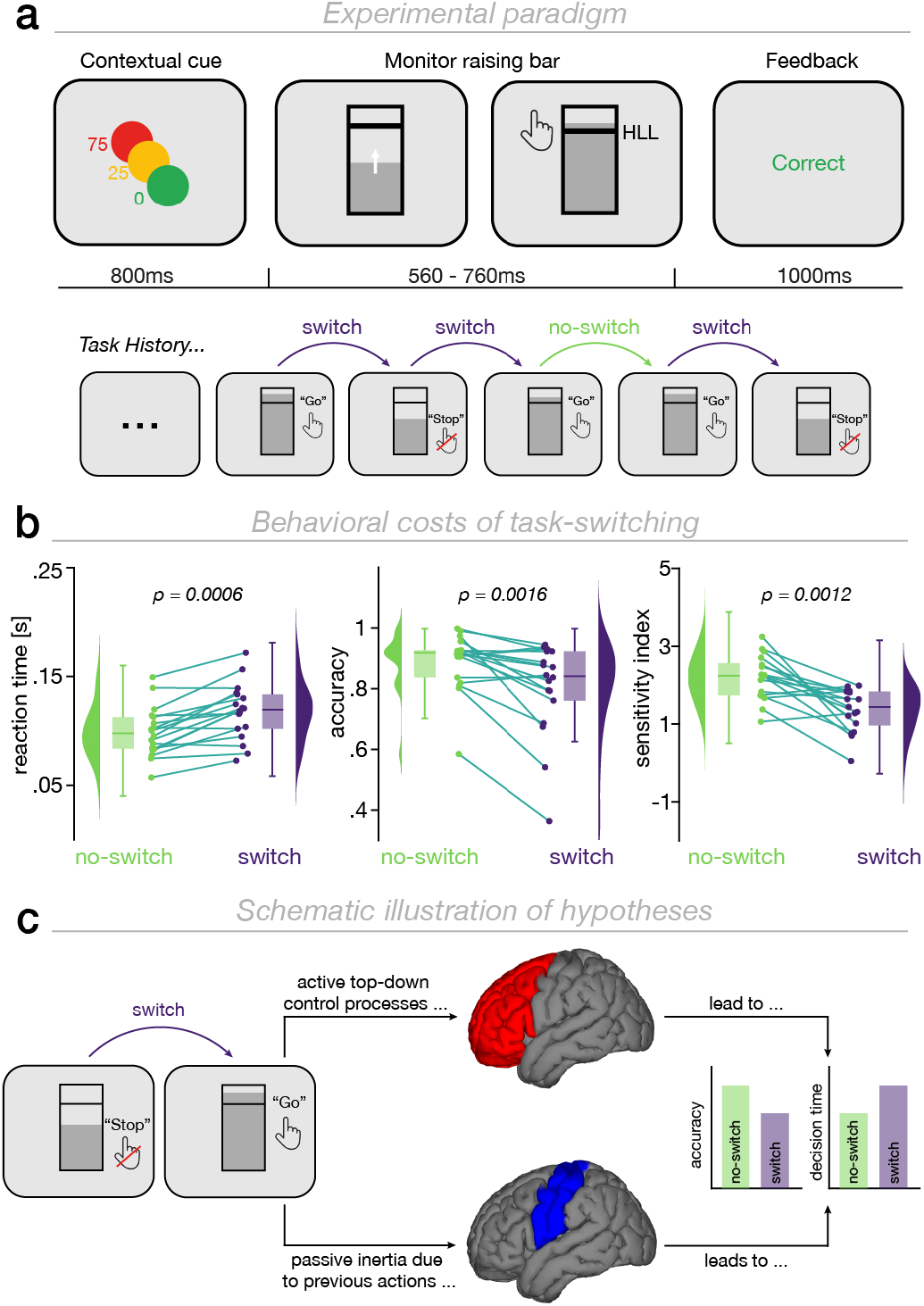
Experimental design, behavioral performance and schematic hypotheses. **a**, *Top:* participants performed a predictive motor task where they had to track a vertically moving target and respond as soon as the target reached a pre-defined lower limit (*go-trials*; hit-lower limit, HLL; black horizontal line). At the start of every trial, participants received a contextual cue indicating the likelihood (0, 25 and 75% likelihood; green, orange, red circle, respectively) that the moving target would stop prematurely requiring participants to withhold their response (*stop-trials*). Feedback was provided at the end of each trial. *Bottom:* task-switching was defined as the trial-type (*stop/go-trial*) congruency between two successive trials (*Methods*). **b**, Behavioral performance as a function of task-switching. *Left:* reaction time was significantly increased after switch trials *(go-trial preceded by a stop-trial)* as compared to no-switch trials *(go-trial preceded by a go-trial)*. Accuracy (*middle*) and *d-prime (right)* significantly decreased after switch as compared to no-switch trials. Grey lines display individual participants, density-plots display the data distribution and boxplots show the median (horizontal line), the first/third quartile (upper/lower edge of box) and the minima/maxima (vertical lines). **c**, Schematic illustration of hypothetical outcomes. In the first scenario, switch costs may reflect the time needed to engage active top-down control processes in the prefrontal cortex to reconfigure the cognitive system. In the second scenario, switch costs could reflect persistent inhibition (passive inertia) of motor areas after withholding a response.

### Shifting between competing task demands has a behavioral switch cost

To assess whether task-switching modulated behavioral performance, we contrasted reaction time, accuracy and d-prime (*d’*) between *switch-* and *no-switch-trials*. Participants were significantly slower (*P* = 0.0006; Cohen’s *d* = 1.29; +19.02 ± 14.73ms; mean ± SD; Wilcoxon rank sum test; **Fig. 1b**) and made significantly more mistakes after *switch-*as compared to *no-switch-trials* (*P* = 0.0016; Cohen’s *d* = -0.9; +8.99 ± 9.94%; mean ± SD**; Fig. 1b**). Furthermore, task-switching across successive trials reduced participants’ sensitivity to correctly decide between two action alternatives (*d’; P* = 0.0012; Cohen’s *d* = -1.04; **Fig. 1b**). We further confirmed that our behavioral analyses were not confounded by regressor collinearity (*Methods;* **Supplementary Fig. 1**). We observed a main effect of *predictive context* and *task-history* on reaction time (*task-history:* 95% CI = [0.016 0.026], *P* < 0.0001; *predictive context:* 95% CI = [0.0005 0.0006], *P* < 0.0001) and accuracy (*task-history:* 95% CI = [-0.99 -0.37], *P* < 0.0001; *predictive context:* 95% CI = [-0.03 -0.02], *P* < 0.0001). Thus, *predictive context* and *task-history* independently modulated behavior.

### Overlapping neural populations encode past and present states in human PFC

To dissect the neural mechanisms underlying behavioral switch costs, we simultaneously recorded neural activity from electrodes located in the human PFC (26 ± 18.74 electrodes per participant; mean ± SD) and motor cortex (14.83 ± 15.23 electrodes). Based on theoretical models^32-34^, two possible scenarios were considered: (1) Switch costs could reflect the time needed to engage active top-down control processes in PFC (**Fig. 1c**) or alternatively, (2) switch costs might result from prolonged inhibition of motor cortex between switching from movement-inhibition to movement-execution (**Fig. 1c**). In order to differentiate these models, we quantified the univariate neural information (unsigned, bias-corrected percent explained variance; *Methods*) about *task-history* and *predictive context* in PFC and motor cortex. We orthogonalized the different factors to disentangle their unique behavioral relevance. We observed significant context- and history-dependent neural information in PFC (**Fig. 2a-d**; *task-history*: *P*_*cluster*_ < 0.0001, Cohen’s *d =* 1.81; *predictive context*: *P*_*cluster*_ < 0.0001, Cohen’s *d* = 2.07; cluster permutation test) and motor cortex (*task-history*: *P*_*cluster*_ < 0.0001, Cohen’s *d* = 1.46; *predictive context*: *P*_*cluster*_ = 0.005, Cohen’s *d* = 2.41).

**Fig. 2.**
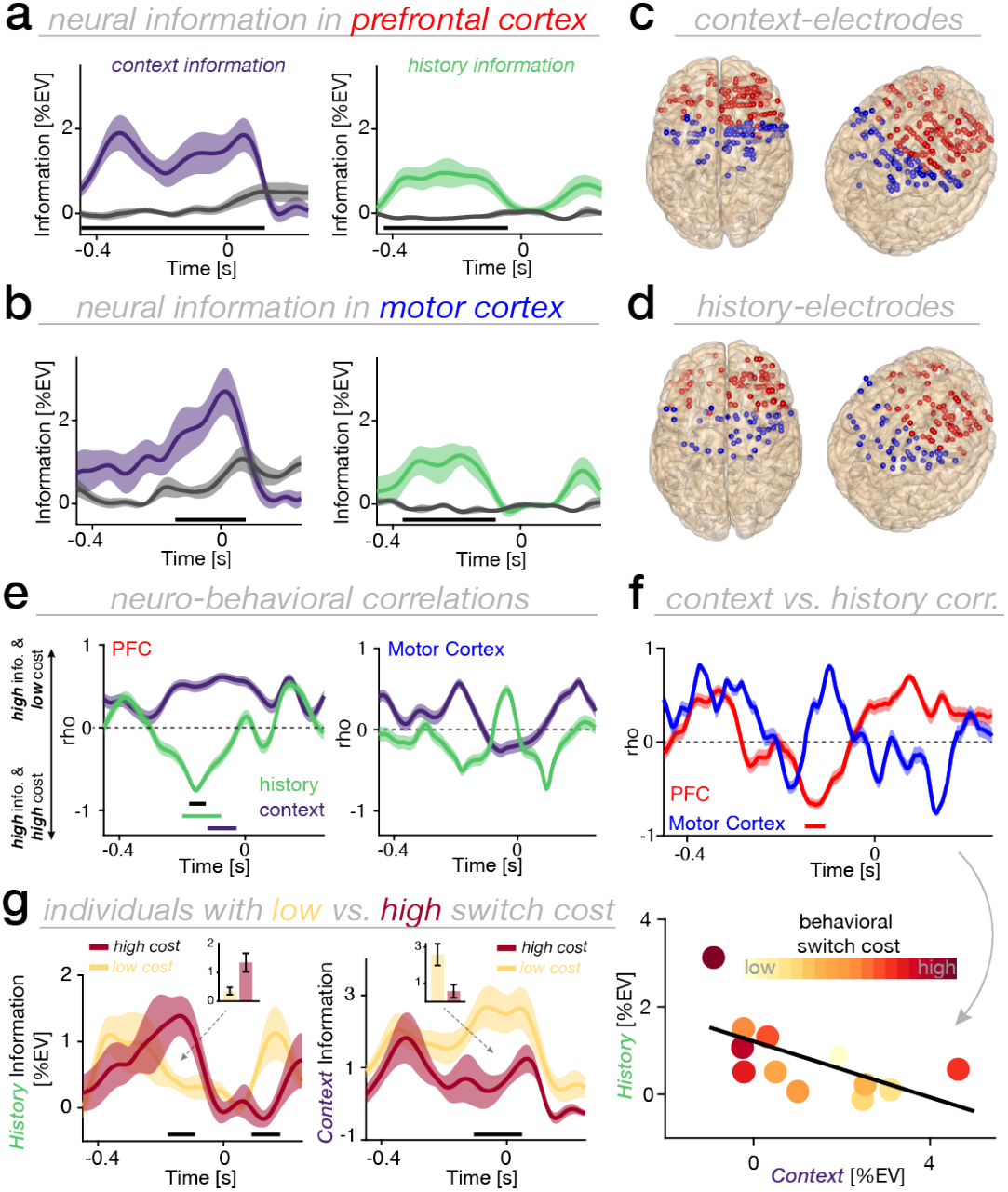
Behavioral dissociation of neural dynamics encoding the past and present. **a**, *Left:* Time course of context-dependent information averaged across all context-encoding electrodes in PFC. *Right:* Time course of history-dependent neural information averaged across all history-encoding electrodes in PFC. Lines and shaded regions show the mean and SEM. Grey traces indicate the time course of neural information across non-encoding electrodes. The lower horizontal black line shows the temporal extent of significant neural information. **b**, Same as **(a)**, but for motor cortex. **c**, Context-encoding electrodes overlaid on a standardized brain in MNI coordinates for PFC (red) and motor cortex (blue). Overall, 165 electrodes in PFC and 89 electrodes in motor cortex carried significant context-dependent information. **d**, History-encoding electrodes, same conventions as in **(c)**. *Task-history* was significantly encoded in 96 electrodes in PFC and 48 electrodes in motor cortex. **e**, *Left:* Temporally resolved correlation between neural information and individual switch costs (accuracy; *see* **Supplementary Fig. 2** for reaction time) for PFC *(left)* and motor cortex *(right)*. Lines and shaded regions show the mean and 95% CIs of bootstrapped correlation coefficients *(Methods)*. The lower colored horizontal lines show the temporal extent of significant correlation for context- (purple) and history-dependent (green) neural information. The black line shows the temporal extent of significant differences in neuro-behavioral correlation between *predictive context* and -history. **f**, *Top:* Temporally resolved correlation between context- and history-dependent neural information for PFC (red) and motor cortex (blue). The red horizontal line shows the temporal extent of significant correlation between context- and history-dependent information in PFC. *Bottom:* correlation between context- and history-dependent information based on the significant temporal cluster shown in the top panel of **(f)**. Filled dots represent individual participants; color-coded by their individual switch costs. **g**, Median split analysis based on individual switch costs (shown for accuracy) for history- *(left)* and context-dependent information *(right)* in PFC. The lower horizontal black line highlights the temporal extent of significant differences between individuals with a low (yellow) versus high (red) switch cost. Lines and shaded regions show the mean and SEM. The small insets depict the averaged neural information across significant clusters for individuals with high versus low switch costs.

Having established a robust coding of *predictive context* and *task-history* in the prefrontal-motor network, we then investigated when neural information coding predicted individual switch costs in behavior using a sliding window correlation analysis (*Methods*). We hypothesized anti-correlated effects of context- and history-dependent neural information on behavior, since an optimal agent should only rely on the currently relevant task context and disregard uninformative features such as *task-history* to efficiently guide decisions.

In line with our hypothesis, strong neural coding of context predicted lower switch costs, whereas strong neural coding of *task-history* indexed increased behavioral costs (**Fig. 2e**; **Supplementary Fig. S2**). Importantly, a significant neuro-behavioral correlation was only observed in PFC (*predictive context*: *P* = 0.039, *r* = 0.59; *task-history*: *P* = 0.013, *r* = -0.57; cluster permutation test), but not in motor cortex (*predictive context*: no cluster; *task-history*: *P* = 0.434). Furthermore, this association differed significantly between *predictive context* and *task-history* for PFC (*P* = 0.028; *motor cortex*: *P* = 0.854; FDR-corrected^35^; black horizontal line in **Fig. 2e**; *Methods*).

Next, we quantified the link between information coding of *predictive context* and *task-history* (**Fig. 2f**; *Methods*), which revealed a robust negative correlation between context- and history-dependent neural information in PFC (*P* = 0.004, *r* = -0.64; cluster permutation test), but not in motor cortex (*P* = 0.227). Note, however, that the correlation between PFC and motor cortex did not significantly differ (*P* = 0.183; *Methods*).

To further illustrate the group-level correlation between information coding and behavioral switch costs, we performed a median split analysis (**Fig. 2g**). Participants with low switch costs revealed stronger neural coding for *predictive context*, but weaker neural coding for *task-history*. The opposite relationship was observed for participants with high switch costs (*predictive context*: *P*_*cluster*_ = 0.026, Cohen’s *d* = -1.6; *task-history*: *P*_*cluster*_ = 0.084, Cohen’s *d* = 1.49; cluster permutation). Collectively, these findings indicate that limited neural resources impose a competition between feature-coding of past and present states and that over-representation of past states comes at substantial behavioral costs.

### Competitive coding of past and present states in human PFC predicts behavior

Having established that neural information coding about past and present states is anti-correlated and exerts dissociable effects on behavior, we tested whether a competitive coding scheme between past and present states in the human PFC could account for individual switch costs. Based on the highly distributed nature of feature-coding in human PFC (cf. **Fig. 2c/d**), a multivariate data analysis approach was used to test this prediction (**Fig. 3a**; *Methods*). We employed a variant of targeted-dimensionality reduction (TDR^36^). In brief, linear regression was used to determine how *predictive context* and *task-history* modulate neural activity at every electrode. Subsequently, low-dimensional subspaces that capture *context*- and *history*-dependent variance in neural activity, were identified using principal component analysis (PCA). Consistent with previous findings demonstrating low-dimensional neural representations of task features^36-38^, we observed that neural coding of *predictive context* and *task-history* was restricted to a low-dimensional subspace (**Fig. 3b**). The activity subspaces in PFC spanned by the first five PCs captured 97.35 ± 0.76% (mean ± SD) of the variance for *predictive context* and 97.33 ± 0.83% of the variance for *task-history* without significant differences in dimensionality between the subspaces (**Fig. 3b**; Wilcoxon rank sum test, *P* = 0.952; comparable for different numbers of components).

**Fig. 3.**
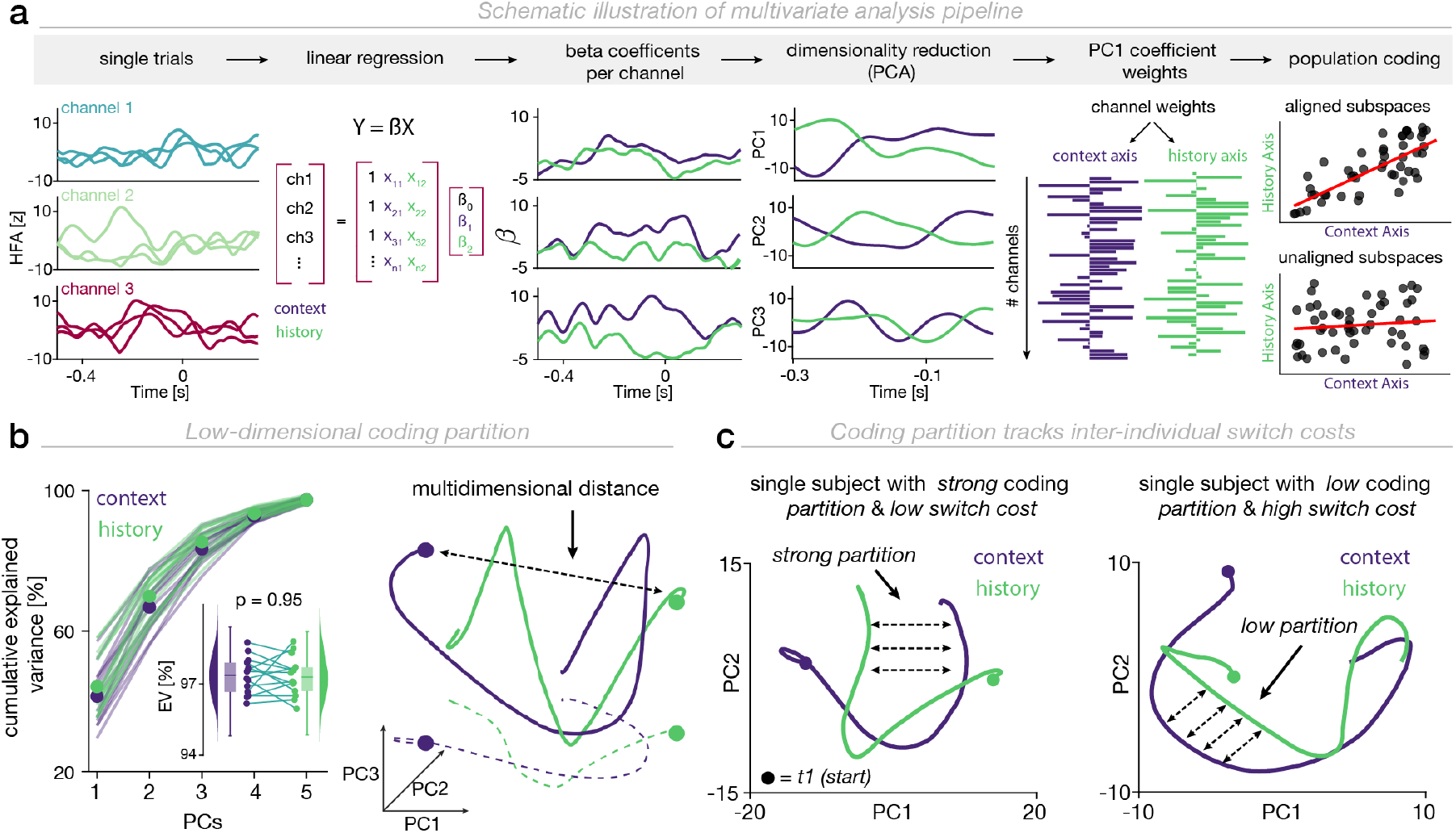
Multivariate data analysis approach to identify low dimensional coding subspaces. **a**, Schematic illustration of the multivariate analysis applied to estimate population coding subspaces *(see Methods)*. **b**, *Left:* Cumulative explained variance estimated using the first five PCs for history- and context-coding subspaces. Small inset illustrates that the dimensionality of context- and history-coding subspaces does not differ. *Right:* Schematic of two coding trajectories through a three-dimensional state space. The black dotted line reflects the multidimensional distance between the two trajectories at time *t = 1*. The colored dotted lines denote the projection line onto PC1 and PC2. **c**, *Left:* Single subject example with a low switch cost and antagonistically evolving context- and history-dependent coding trajectories projected onto the first two PCs. *Right:* Single subject example with a high switch cost and strongly resembling coding trajectories projected onto the first two PCs (*see* **Supplementary Fig. 3** for group-level correlation)

Activity within these subspaces reflects time-varying neural population dynamics predictive of past and present states. Based on the principle of communication subspaces^39^, we tested whether the magnitude of partitioned information between the feature-subspaces predicts individual switch costs. Therefore, we computed the time-resolved multidimensional distance between the two coding trajectories (projected into a common subspace; *Methods*) and extracted the magnitude of maximal divergence between the population coding trajectories (**Fig. 3b** right; *Methods*). In line with our hypothesis, a stronger coding subspace partition (less overlap between coding subspaces) predicted reduced switch costs (**Fig. 3c**; *P* = 0.015, Spearman *r* = -0.64, *N* = 14; **Supplementary Fig. 3**).

### Flexible change in population geometry reduces switch costs

What might be a computational mechanism that enables neural ensembles to partition distinct coding features and thereby allows downstream readers to robustly decode feature-specific information? A viable mechanism would be a flexible, feature-dependent set of weights between upstream and downstream neural populations supporting robust read-out of feature-specific information. To test this prediction, we quantified the *subspace alignment* (**Fig. 3a**; *Methods*) between the coding subspaces accounting for variance of *past* and *present task-features*. A positively aligned subspace indicates that neural ensembles share the same low-dimensional population code for *past* and *present* states (e.g. similar local populations contributing to the respective coding). Consequently, downstream populations would not be able to untangle information about past and present states, ultimately leading to information interference between them. In line with this proposed mechanism, we observed that a stronger alignment between subspaces representing past and present states comes with a stronger individual switch cost (*P* = 0.002, Spearman *r* = 0.78, *N* = 14; **Fig. 4a-c**). Taken together, this set of observations suggests that conjunctive coding of the past and present leads to mutual interference between the past and present in the human PFC, ultimately deteriorating behavioral performance; whereas, efficient coding in distinct population subspaces benefits behavior and enables rapid switching and might therefore, constitute a central mechanism underlying cognitive flexibility.

**Fig. 4.**
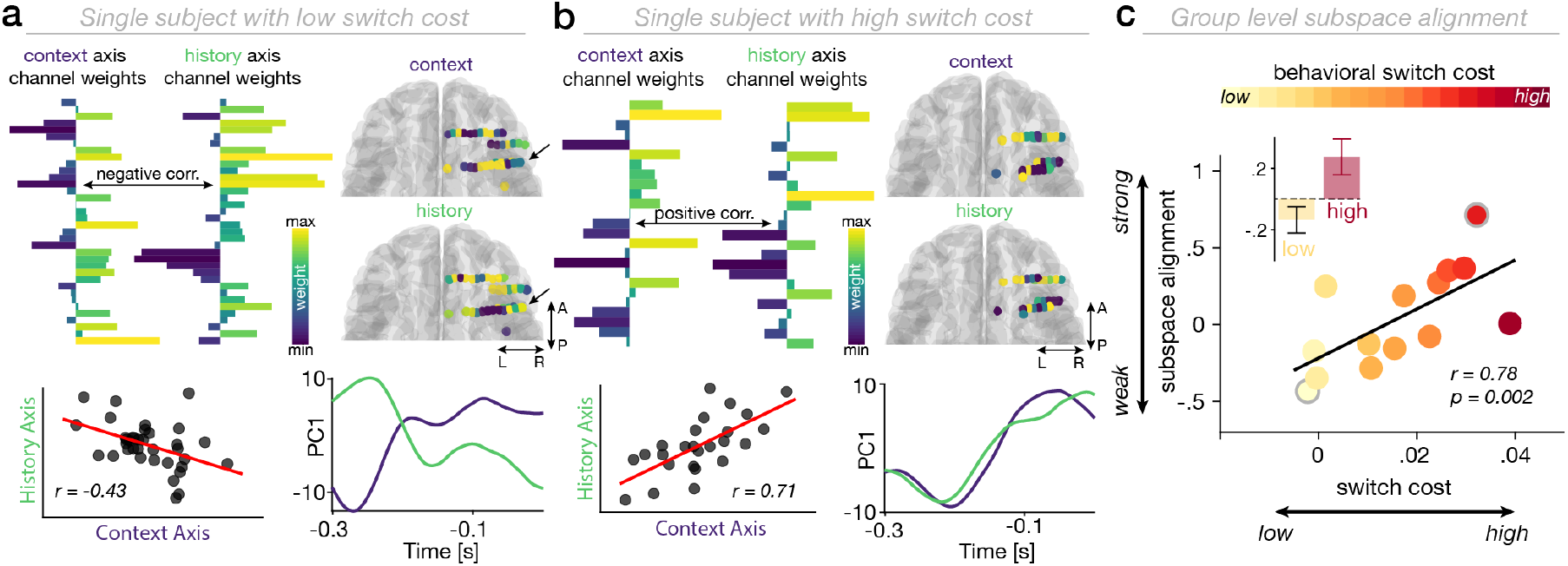
Low-dimensional coding partition predicts individual switch costs. **a**, Example of subspace alignment in PFC for a participant with a low behavioral switch cost. *Top Left:* Channel weights obtained for the dominant mode (PC1) across context *(left)* and history axes *(right). Top Right:* Channel weights obtained for the dominant mode across context *(top)* and history axes *(bottom)* overlaid on an individual brain. *Bottom Left:* Strong negative correlation between channel weights projecting onto the dominant mode across context and history axes. *Bottom Right:* Temporal evolution of low-dimensional context- and history-coding subspaces, shown for the dominant mode. Note the anticorrelated traces. **b**, Example of subspace alignment in PFC for a participant with low behavioral switch costs. Same conventions as in **(a). c**, Group-level correlation reveals a positive relationship between subspace alignment strength and individual switch cost. Strong subspace alignment is associated with a high switch cost whereas weak subspace alignment is associated with a low switch cost. Outlined circles in grey indicate the example subjects in **(a)** and **(b)**. The inset highlights the strength of subspace alignment based on a median split for participants with a low (yellow) and high (red) switch cost.

## Discussion

Humans rapidly switch between actions to meet shifting task demands or internal goals^40^. Flexibly shifting between competing task demands incurs a *switch cost* reflected in slower and more error-prone behavioral responses^21,41^. Despite robust behavioral evidence of switch costs^22,41-43^, their underlying neural mechanisms and neurocomputational principles to overcome interference and diminish switch costs remain elusive. Prior studies have implicated various regions in switching, including prefrontal^1,32,42,44,45^ and sensorimotor regions^46-49^. Yet, the majority of evidence was obtained from single unit recordings or local field potentials in rodents^45^ or non-human primates^1,32,50-52^. Thus, it remains unclear if similar principles apply to the human brain. Here, we bridged this gap using intracranial recordings from the human prefrontal and motor cortex. Our findings demonstrate that (I) neural coding of past and present states coexists in the human prefrontal and motor cortex (**Fig. 2a-d**), and that (II) coding competition between past and present states manifests in switch costs (**Fig. 2e-g**). Crucially, (III) the human PFC solves the interference problem by efficiently partitioning information about the past and present into distinct low-dimensional neural states (**Fig. 3c; Fig. 4a-b**). Finally, (IV) inter-individual variability of low-dimensional coding partition predicts inter-individual variability in switch costs (**Fig. 4c**). In sum, these findings reveal a fundamental coding principle that might constitute a central building block of human cognitive flexibility.

### Neural coding of past and present behavioral states in human PFC and motor cortex

Prior studies have demonstrated that task-relevant features are encoded in a distributed network of brain regions, including frontoparietal^36,53-55^ and sensorimotor regions^56-58^. Recently, it had been shown that the conjunctive neural representation of goal-relevant features is critical for action selection in humans^10^. However, despite mounting evidence emphasizing the strong influence that past states can exert over goal-relevant present states *(serial biases)*, little is known about the intricate interplay between past and present state representations in the human brain. Using direct brain recordings in humans, we demonstrate that neural information about the past and present co-exist in space and time in the human prefrontal and motor cortex. Moreover, univariate information coding revealed similar temporal profiles distributed across both regions. These results go substantially beyond recent evidence obtained in animal models^26^ and show that neural information about the past and present is not unique to the PFC, but is equally represented in the human motor cortex. In sum, these results demonstrate a widely distributed coding of past and present states across the prefrontal-motor hierarchy.

### Competition among past and present states in the human PFC manifest in switch costs

When the past does not predict the future, an optimal agent should in principle discard irrelevant information about the past in order to efficiently guide currently relevant decisions. Prior studies across different species have identified several neural signatures that correlated with past events and choice history during perceptual decision-making, highlighting the key roles of prefrontal^59,60^, parietal^18,61,62^ or motor^18,63^ cortex in shaping subsequent behavior. Most previous studies focused on neural signatures of past states without accounting for co-emerging neural signatures of present states. This precluded strong inference about how co-existing neural dynamics of past and present states jointly shape behavior. Here, we overcome this limitation by combining human iEEG recordings with information theoretical approaches to quantify the neural information about past and present states directly from HFA as a proxy of multi-unit activity^27,28,64^. We demonstrate a behaviorally relevant dissociation between neural coding of past and present states. Specifically, the strength of neural coding of past and present information predicts individual switch costs. Strong neural coding of the present and weak neural coding of the past predicts low switch costs whereas the opposite pattern increases switch costs. These results suggest that limited neural resources give rise to a biased competition between co-existing representations of past and present states: An over-representation of the past interferes with goal-directed behavior in the present and is detrimental for behavior.

### Efficient low-dimensional coding partition of past and present mitigates switch costs

Is there a neural mechanism that reduces mutual interference between competing representations? Theoretical work has proposed that neural populations might resolve interference by orthogonalizing competing representations^23,24^. In this way, the same neural population encodes a competing set of stimuli, but keeps their representations separable in neural state space. This enables downstream populations to optimally decode information about a particular state. Evidence supporting this notion has been obtained in two recent animal model studies^25,26^. These studies demonstrated that both sensory (auditory cortex^25^) and association regions (medial PFC^26^) can maintain competing internal or external inputs by generating orthogonal representations. Yet, whether these representations have immediate behavioral relevance remained unaddressed. Here, our results reveal that the human PFC efficiently resolves mutual interference by partitioning information about past and present states into distinct low-dimensional subspaces. Importantly, the magnitude of overlap between past and present states predicts inter-individual switch costs. These results provide evidence that the segregation of competing representations into distinct population subspaces delineates an efficient coding mechanism that has immediate behavioral relevance constituting an integral component of flexible cognitive control.

### Conclusion

Collectively, these findings uncover a fundamental computational principle how the human PFC resolves interference between competing past and present information. The results establish that competition between overlapping representations of past and present states can be reduced by partitioning state-specific information into non-overlapping, low-dimensional coding subspaces.

## Materials and Methods

### Participants

We obtained intracranial recordings from a total of 19 pharmaco-resistant epilepsy patients (33.73 years ± 12.52, mean ± SD; 7 females) who underwent presurgical monitoring and were implanted with intracranial depth electrodes (DIXI Medical, France). Data from one patient were excluded from neural analyses because a low-pass filter was applied at 50 Hz during data export from the clinical system, thus, precluding analyses focusing on high-frequency activity. All patients were recruited from the Department of Neurosurgery, Oslo University Hospital. Electrode implantation site was solely determined by clinical considerations and all patients provided written informed consent to participate in the study. All procedures were approved by the Regional Committees for Medical and Health Research Ethics, Region North Norway (#2015/175) and the Data Protection Officer at Oslo University Hospital as well as the University Medical Center Tuebingen (049/2020BO2) and conducted in accordance with the Declaration of Helsinki.

### iEEG Data Acquisition

Intracranial EEG data were acquired at Oslo University Hospital at a sampling frequency of 512 Hz using the NicoletOne (Nicolet, Natus Neurology Inc., USA) or at a sampling frequency of 16 KHz using the ATLAS (Neuralynx) recording system.

### CT and MRI Data Acquisition

We obtained anonymized postoperative CT scans and pre-surgical MRI scans, which were routinely acquired during clinical care.

### Electrode Localization

Two independent neurologists visually determined all electrode positions based on individual scans in native space. For further visualization, we reconstructed the electrode positions as outlined recently^65^. In brief, the pre-implant MRI and the post-implant CT were transformed into Talairach space. Then we segmented the MRI using Freesurfer 5.3.0^66^ and co-registered the T1 to the CT. 3D electrode coordinates were determined using the Fieldtrip toolbox^67^ on the CT scan. Then we warped the aligned electrodes onto a template brain in MNI space for group-level analyses.

### Experimental Procedure

Participants performed a predictive motor task where they had to continuously track a vertically moving target and respond as soon as the target hits or withhold their response if the target stops prior a predefined spatial position using their dominant hand^30^. Each trial started with a baseline period of 500ms followed by a cue (presented for 800ms centered) that informed participants about the likelihood that the moving target would stop prior to the lower limit (hit lower limit; HLL; **Fig. 1a**). Thus, the predictive cue could be directly translated into the probability that participants had to release the space button (BR) or withhold their response. Participants were instructed to either release the button as soon as the target hits (*go-trials*) or withhold their response if the target stops prior to the HLL (*stop-trials*). We parametrically modulated the likelihood of stopping. A green circle indicated a 0% likelihood, an orange circle indicated a 25% likelihood and a red circle indicated a 75% likelihood that the moving target would stop prior to the HLL. Upon receiving the predictive cue, participants were able to start the trial in a self-paced manner by pressing the space bar on the keyboard. By pressing the space bar, the target would start moving upwards with constant velocity and reach the HLL after 560 – 580ms. The upper boundary was reached after 740 – 760ms, thus, leaving a response window of 160ms between the HLL and the upper boundary. If participants released the button within this 160ms interval, the trial was considered as correct. Trials in which the button was released either before or after this interval were considered as incorrect. Feedback was provided after every trial for 1000ms.

### Behavioral Analysis

The effect of *predictive context* on behavioral performance has been extensively reported in a previous study^30^. Here, we defined task-switching based on the congruency between the trial type (*stop/go-trial*) at trial *N* and at trial *N-1* (*congruent = go-trial followed by go-trial; incongruent = stop-trial followed by go-trial*). Note that, given the experimental setup, the reverse step (*go-trial followed by stop-trial*) was not feasible since our analyses required time-locking relative to the motor response, which was withheld in case of stop-trials. To account for collinearity between various factors (**Supplementary Fig. 1**) on behavioral performance, we employed a generalized linear mixed effect model including *current predictive context (likelihood of stop), past predictive context (likelihood of stop in past trial), task-history (stop/go-trial), past choice (button release/withhold response)* and *past feedback (correct/incorrect)* as fixed-effect predictors, *participants* as random-effects and *response time/accuracy* as response variables.

### iEEG Preprocessing

Intracranial EEG data were demeaned, linearly de-trended, locally re-referenced (bipolar derivations to the next adjacent lateral contact) and if necessary down-sampled to 512 Hz. To remove line noise, data were notch-filtered at 50 Hz and all harmonics. Subsequently, a neurologist visually inspected the raw data for epileptic activity. Channels or epochs with interictal epileptic discharges (IEDs) and other artifacts were removed. We segmented the cleaned data into 10 seconds long, partially overlapping trials to prevent edge artifacts due to subsequent filtering. Unless stated otherwise, trials were event locked to the participants’ response.

### Extraction of High-Frequency Activity (HFA)

The extraction of the high-frequency activity time series was conducted in a three-step process. First, we bandpass-filtered the raw data epochs (10 seconds) between 70-150 Hz into eight, non-overlapping 10 Hz wide bins. We then applied the Hilbert transform to obtain the instantaneous amplitude of the filtered time series. In a last step, we normalized the high-frequency traces using a bootstrapped baseline distribution^68,69^. This involved randomly resampling baseline values (from -0.2 to -0.01s relative to cue onset) 1000 times with replacement and normalizing single high-frequency traces by subtracting the mean and dividing by the standard deviation of the bootstrap distribution. The eight individual high-frequency traces were then averaged to yield a single time series of high-frequency band activity.

### Regions of Interest

We classified electrodes into discrete prefrontal or (pre-)motor electrodes based on anatomical and functional characteristics using the human Brainnetome Atlas^70^ (*see* **Supplementary Table 1** for details about electrode classification).

### Univariate Information Dynamics

We quantified the information encoded in a neural population about two main factors of interest, *predictive context* and *task-history*, using a well-established information theoretical approach^55,71-73^. We employed a six-way analysis of variance (ANOVA) to quantify the percentage of HFA variance explained by the following task factors: *predictive context (likelihood of stop), choice, past predictive context (likelihood of stop in last trial), task-history (stop or go trial), past choice, past feedback*. Importantly, due to collinearity between the task factors, an unbalanced ANOVA^55^, that implicitly orthogonalized the different factors, was employed. Thus, variance explained by *task-history* or *predictive context* could not be explained by any other residual regressor. The amount of percent explained variance was quantified using the debiased effect size ω^2^, which is defined as

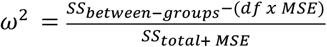

where *SS*_*total*_ reflects the total sum of squares across *n* trials,

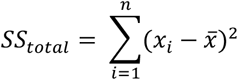

*SS*_*between–groups*_ the sum of squares between *G* groups (e.g. factor levels),

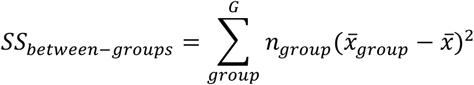

*MSE* the mean square error,

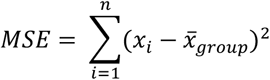

and *df* the degrees of freedom specified as *df*=*G*−1. We estimated ω^2^ using a sliding window of 50ms that was shifted in steps of 2ms to obtain a time course of neural information. This approach is insensitive with respect to the time of task-related activation and to the direction of encoding (i.e., HFA increases or decreases). Electrodes that revealed a significant main effect of *task-history* and/or *predictive context* for at least 10% of the trial length were considered as information-encoding electrodes^68,71,74,75^. Finally, to minimize inter-individual variance and maximize the sensitivity to identify a temporally consistent pattern that accounts for most of the variance across participants, we used principal component analysis (PCA)^72,75^. PCA was applied to the *F* value time series concatenated across participants (channel x time matrix^30,72,76^). In order to define PCs that explain a significant proportion of variance in the data, we employed non-parametric permutation testing (1000 iterations) to determine the proportion of variance explained by chance. Electrodes that exhibited a strong weight (75^th^ percentile) on any of the high variance-explaining PCs were used for further analyses as outlined recently^72^.

The orthogonalized computation of neural information allowed us to further quantify the similarities of information dynamics linked to *predictive context* and *task-history* in a time-resolved manner using spearman correlation. To further link the strength of regressor information to the individual switch costs, we first computed the individual switch cost by subtracting reaction time/accuracy on *no-switch-trials* from reaction time/accuracy on *switch-trials*. In a second step, we then correlated the individual switch cost with the strength of neural information related to *predictive context* or *task-history* in a time-resolved manner. Significance of correlation across time was assessed using cluster-based permutation testing to correct for multiple comparisons (1000 iterations; randomly shuffling participant indices without replacement). To compare two distributions of correlation coefficients, we used Fisher’s z-transformation to convert Pearson’s *r* to the normally distributed variable *z*, based on which the *p-value* was derived.

### Multivariate Population Dynamics

We characterized low-dimensional coding dynamics with respect to *predictive context* and *task-history* using a variant of targeted-dimensionality reduction (TDR^36^). A multi-variable linear regression was employed to determine how *predictive context* and *task-history* contribute to the response of every electrode in a temporally resolved manner. Here, we included all electrodes with no prior constraint on information strength to estimate latent coding dynamics across the entire sampled population. Subsequently, principal component analysis (PCA) was employed to identify low-dimensional subspaces capturing variance due to *predictive context* and *task-history*. The subsequent population analysis focused on the time window prior to the participants’ response (300ms to 0ms; cf. **Fig 2e/f**) to maximize the temporal sensitivity of population coding analysis.

#### Neural coding trajectories

To verify that coding trajectories are low-dimensional, we computed the cumulative explained variance of the first 5 principal components (PCs). We then quantified the Euclidean distance *D* between coding trajectories for *predictive context* and *task-history* in a temporally resolved manner (window size = 50ms; shift size = 10ms) as

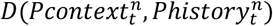

where *P* represents a point in the context- or history-trajectory embedded a n-dimensional space where *n* reflects the number of PCs required to cumulatively explain at least 95% of the total variance. Importantly, we first projected the *history-subspace* into the *context-subspace*^77,78^. We then determined the magnitude of maximal divergence between the two coding trajectories across time to further infer the behavioral relevance of coding separation between *predictive context* and *task-history* on a group level. We excluded participants with less than 10 electrodes per ROI from this analysis to ensure robust PCA estimates.

#### Subspace alignment analysis

In a final step, we computed the alignment of low-dimensional coding subspaces. We used the coefficient matrix based on the PCA approach outlined above which contains the individual channel weights per principal component. We quantified alignment *A* between low-dimensional context- and history-coding subspaces as the correlation coefficient (Spearman’s Rho) between vector *C* containing the weights of all electrodes *n* contributing to the subspace defined by *predictive context* and vector *H* containing the weights of all electrodes *n* contributing to the subspace defined by *task-history A = corr*(*C*_*n*_, *H*_*n*_). We excluded participants with less than 10 electrodes per ROI from this analysis.

### Statistics

Unless stated otherwise, we used non-parametric cluster-based permutation testing^79^ to analyze data in the time domain (**Fig. 2a/b/e/f**).

#### Neural information

Clusters for neural information time series (**Fig. 2a/b/g**) were formed by thresholding a dependent (**Fig. 2a/b**) or independent (**Fig. 2g**) t-test at a critical alpha of 0.05. We generated a permutation distribution by randomly shuffling encoding vs. non-encoding electrode labels (**Fig. 2a/b**) or condition labels (**Fig. 2g**) and recomputing the cluster statistic. The permutation p-value was obtained by comparing the cluster statistic to the random permutation distribution. Clusters were considered to be significant at *P* < 0.05.

#### Correlation analyses

Clusters for correlation time series (**Fig. 2e/f**) were formed by thresholding the resulting correlation p-values at 0.05. We generated a permutation distribution by randomly shuffling participant labels and recomputing the cluster statistic. The permutation p-value was obtained by comparing the cluster statistic to the random permutation distribution. Clusters were considered to be significant at *P* < 0.05.

## Acknowledgments

We thank Anais Llorens and Ingrid Funderud for their help with data collection. This work was funded by the Baden Wuerttemberg Foundation (Postdoc Fellowship; RFH), German Research Foundation, Emmy Noether Program (DFG HE8329/2-1; RFH), Hertie Foundation, Network for Excellence in Clinical Neuroscience (RFH), the International Max Planck Research School for the Mechanisms of Mental Function and Dysfunction (GI), the Research Council of Norway (grant number 240389; AKS, TE, PGL; grant number: 314925; AOB), the Research Council of Norway (Centre of Excellence scheme, grant number 262762; RITMO, RITPART International Partnerships for RITMO Centre of Excellence, grant number 274996; AKS, TE, PGL) and by a NIMH Conte Center Grant (1 PO MH109429, RTK) and the NINDS (2 R01 NS021135, RTK).

## Author contributions

Conceptualization: JW, GI, RFH

Methodology: JW, GI

Investigation: AKS, TE, PL, JI, AOB

Visualization: JW, GI

Funding acquisition: AKS, RTK, TE, RFH

Project administration: TE, AKS, RFH

Supervision: RFH

Writing – original draft: JW, RFH

Writing – review & editing: GI, AKS, RTK, TE, AOB, JI, PL

## Data availability

Source data is included as Supporting Information. Raw data are available upon request from Anne-Kristin Solbakk or Tor Endestad.

## Code availability

Freely available software and algorithms used for analysis are listed where applicable. All code will be made publicly available upon publication on GitHub.

